# Ccn2 deficiency causes smooth muscle cell de-differentiation and severe atherosclerosis in hyperlipidemic mice

**DOI:** 10.1101/2023.01.16.524161

**Authors:** JH Larsen, J Stubbe, HC Beck, M Overgaard, CA Lindegaard, DR Hansen, R Goldschmeding, RR Rodriguez-Díez, M Ruiz-Ortega, C Pyke, RC Wirka, JS Lindholt, LM Rasmussen, LB Steffensen

**Author notes:** **CORRESPONDING AUTHOR** Lasse Bach Steffensen, J. B. Winsløws Vej 21, 3-22A, 5000 Odense C, Denmark, Mail.

## Abstract

Cellular communication network factor 2 (CCN2/CTGF) is a matricellular protein with an established role in fibrotic diseases and cancers, and therapies targeting CCN2 is currently in Phase II and III clinical trials for idiopathic pulmonary fibrosis, pancreatic cancer and Duchenne Muscular Dystrophy. Recent studies have highlighed a protective role of CCN2 in aortic aneurysm disease, but its role in atherosclerosis remains to be investigated.

We identified arteries as having the highest relative expression of *CCN2* across 54 human tissues. In aortas, *CCN2* was among the highest expressed genes, and *in situ* hybridization of human internal thoracic arteries revealed vascular smooth muscle cells (SMCs) as its principal source.

Hypothesizing a role for CCN2 in SMC phenotype maintenance and athero-protection, we investigated inducible Ccn2 knockout (*Ccn2*^*Δ/Δ*^) mice in normo- and hyper-lipidemic settings. Induction of hyperlipidemia by single intravenous injection of 1·10^11^ viral genomes of rAAV8-D377Y-mPcsk9 combined with 24 weeks of western type diet resulted in severe enlargement (3-5-fold increase of relative aorta mass compared to wildtype littermates, *p* < 0.0001) and whitening of *Ccn2*^*Δ/Δ*^ aortas. Oil Red O-staining of *en face* prepared thoracic aortas showed a marked increase in atherosclerosis in *Ccn2*^*Δ/Δ*^ mice as compared to wildtype littermates (75% vs. 10% Oil Red O-positive aortic area, *p* < 0.0001). Transcriptomic profiling of cultivated SMCs derived from aortas of normolipidemic mice showed signatures of dedifferentiation (reduced expression of *e*.*g. Myocd, Acta2* and *Myh11*) and modulation toward a synthetic, pro-inflammatory phenotype of *Ccn2*^*Δ/Δ*^ SMCs. These effects were verified *in vivo* and in *CCN2*-silenced human aortic SMCs. Taken together, we find that CCN2 plays a critical athero-protective role in artery tissues, likely through maintaining SMCs in a differentiated, contractile phenotype.

## INTRODUCTION

Vascular smooth muscle cells (SMCs) constitute the bulk of artery tissues. These highly specialized cells are the principal producers of structural extracellular matrix (ECM) of the artery wall, and they are essential for regulating vascular tone(1). Fully differentiated SMCs residing in the medial layer of arteries are rich in intracellular myofilaments. Myocardin (MYOCD) is regarded the guardian of the contractile SMC. It is a co-activator of Serum response factor (SRF), which binds CArG elements in promoter regions of virtually all genes encoding contraction-related proteins(2). During atherogenesis, environmental stimuli cause downregulation of MYOCD expression and/or activity in local SMCs and consequently downregulation of myofilaments(3). The concept of phenotypic modulation of contractile SMCs and the involvement of SMC-derived cells in atherosclerosis has long been acknowledged(4, 5), and SMCs invading the intimal layer have been characterized as “synthetic” having abundant secretory organelles, *i*.*e*. rough endoplasmatic reticulum (ER) and Golgi-apparatus(6). Reliance on traditional SMC markers (*e*.*g*. ⍺ smooth muscle actin (ACTA2) or smooth muscle cell myosin heavy chain (MYH11)) has traditionally ascribed SMCs a beneficial role in atherosclerosis, since ACTA2^+^/MYH11^+^ plaque-cells are located in the fibrous cap and likely the principal producers of the plaque-stabilizing ECM(7). In more recent years, this perception has been challenged. Today, the emerging use of mono- and multicolor lineage-tracer mouse models have unambiguously unveiled that SMC-derived plaque cells originate from few medial SMCs(8, 9), that they contribute to 30-70% of plaque cells(8-13), and that they can adopt a wide spectrum of plaque phenotypes(3, 9, 11, 14). While these recent technological advancements have established the importance of SMCs in atherogenesis, current efforts aim at identifying factors governing SMC fate in atherosclerosis, as such insight may disclose new therapeutic options to promote stabilization of plaques in order to prevent lethal rupture(1).

Cellular communication network factor 2 (CCN2, previously known as Connective tissue growth factor (CTGF)(15)) is a 38 kDa matricellular protein, which has been extensively studied in the context of fibrosis and cancer, in which CCN2 expression is markedly elevated(16). An important role for CCN2 in artery tissues has emerged in recent years, as Ccn2 depletion augments aortic aneurysm development in mouse models of this disease(17, 18).

CCN2 is not a growth factor as indicated by its original name, and no specific CCN2 receptor has been identified. Rather, CCN2 is a multidomain protein capable of binding various integrins, non-integrin receptors, ECM components, cytokines, growth factors and proteases, and the myriad of cellular effects attributed to CCN2 appear to be highly tissue- and context-dependent(16).

CCN2 has emerged as a promising target in various fibrotic conditions, and FG-3019/Pamrevlumab, an anti-CCN2 antibody, is currently in Phase II and III clinical trials for idiopathic pulmonary fibrosis (IPF), pancreatic cancer and Duchenne Muscular Dystrophy (ClinicalTrials-gov identifiers: NCT03955146, NCT03941093, NCT02606136).

Here, we report that *CCN2* is one of the highest expressed genes in healthy artery tissues, and we demonstrate a critical role of CCN2 in the maintenance of artery integrity and protection against atherosclerosis. These findings merit caution in using anti-CCN2-based therapy.

## RESULTS

### CCN2 is among the highest expressed genes in healthy artery tissues

The gene-tissue expression (GTEx) portal(19) was used to assess *CCN2* transcript levels across 54 human tissues. Relative *CCN2* expression was elevated in artery tissues as compared to all other tissues (**Figure 1A**). In the aorta, *CCN2* was among the 30 genes with highest expression (excluding housekeeping- and mitochondrial genes) (**Figure 1B**), and in internal thoracic artery (ITA) specimens (*n* = 4) obtained from coronary artery bypass grafting (CABG), *CCN2* was readily detectable and identified primarily in vascular smooth muscle cells (SMCs) in the medial layer (**Figure 1C-D**).

**Figure 1.**
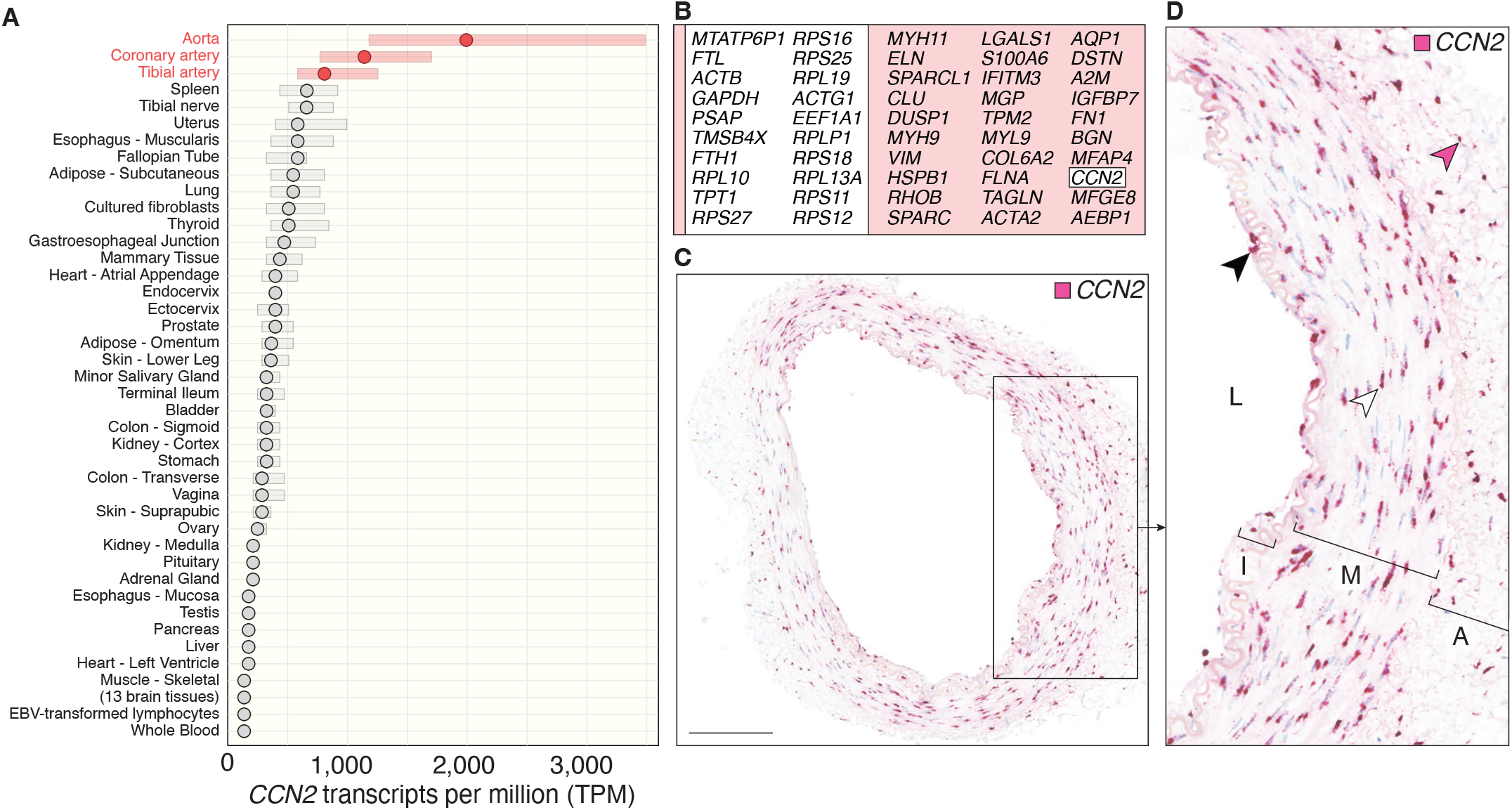
CCN2 is among the highest expressed genes in artery tissues. **A**. *CCN2* transcripts per million (TPM) across all tissues from the GTEx portal represented as median (dots) and interquartile range (bars). The three artery tissues (ascending aorta (*n* = 432), noncalcified coronary artery (right and left) (*n* = 240) and left tibial artery from the gastrocnemius region (*n* = 663)) are shown in red. **B**. The 50 non-mitochondrial genes with highest TPM in aorta according to the GTEx portal. Genes in large white box are highly expressed in most tissues and can be considered housekeeping genes. **C**. *CCN2* ISH of human ITAs. Scalebar = 250 *μ*m. **D**. Magnification of rectangle in C. White, black and pink arrowheads point to *CCN2*-positive SMC-, endothelial- and adventitial cells, respectively. L: Lumen, I: Intima, M: Media, A: Adventitia.

### Ccn2 deficiency causes structural abnormalities of the aorta

Hypothesizing a critical role of CCN2 in artery tissue, we turned to a mouse model of Ccn2 deficiency. Constitutive *Ccn2* knockout (*Ccn2*^*-/-*^) mice die shortly after birth due to respiratory failure(20). We therefore used a tamoxifen-inducible *Ccn2* knockout mouse model. All mice were homozygous for *loxp*-flanked *Ccn2* (*flCcn2*), while only inducible knockout mice harbored *Rosa26-CreER*^*T2*^. Six weeks post tamoxifen administration, the amplicon representing *flCcn2* was undetectable in tail biopsies (**Figure 2A**), aortic *Ccn2* mRNA was abolished, and aortic Ccn2 protein was reduced to 20% in *Cre*-positive mice (henceforth referred to as *Ccn2*^*Δ/Δ*^ mice) (**Figure 2B-E**). Wildtype (*Ccn2*^*+/+*^) and *Ccn2*^*Δ/Δ*^ mice were similar in weight (**Figure 2F**), but female *Ccn2*^*Δ/Δ*^ mice had reduced relative heart mass (**Figure 2G**). Histological assessment of thoracic aortas revealed a morphologically disorganized aortic wall with increased media cross-sectional area (CSA) (**Figure 2H-I**).

**Figure 2.**
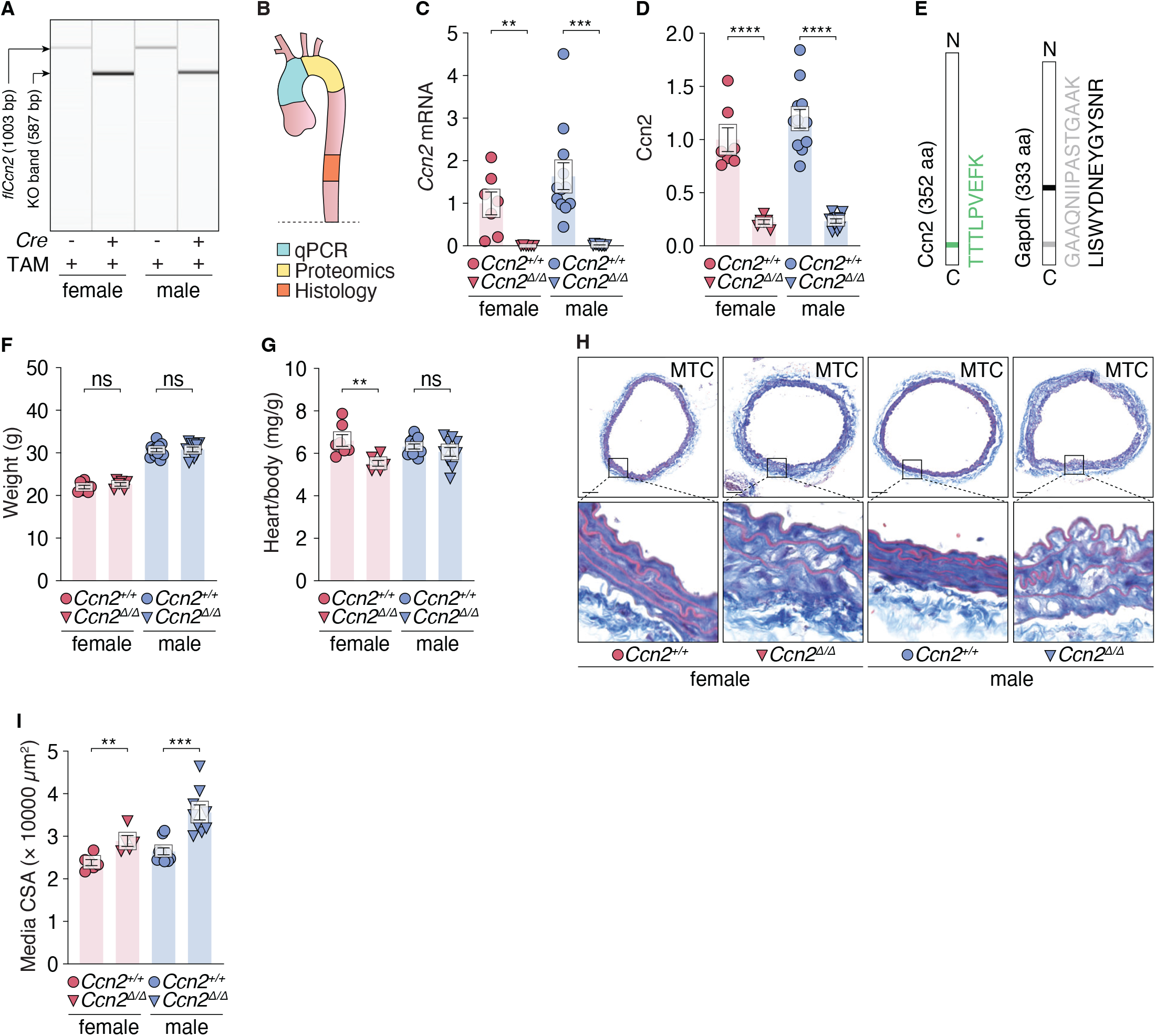
Ccn2 deficiency causes severe structural abnormalities of the aorta. **A**. Representative genotyping results of tamoxifen-treated *flCcn2* mice with and without *Cre*. **B**. Aorta segments used for qPCR (C), proteomic (D-E), and histological (H-I) analysis. Dotted line represents the diaphragm. **C**. *Ccn2* mRNA expression in the aorta. **D**. Ccn2 protein level in the aorta. For C-D, data are normalized to the mean of female *Ccn2*^*+/+*^ mice. **E**. Peptides of Ccn2 and Gapdh used in the MRM assay used to obtain results in D. The position of peptides is indicated. aa = amino acids. **F-G**. Weight (F) and relative heart mass (G) of normolipidemic mice. **H**. Representative cross-sections of aortas stained by MTC. Scalebars = 100 *μ*m. **I**. Quantitation of aorta media CSA. Barplots in the figure show mean ± SEM. *: *p* ≤ 0.05, **: *p* ≤ 0.01, ***: *p* ≤ 0.001, ****: *p* ≤ 0.0001, ns: non-significant. Unpaired t-test with Welch’s correction was used in C-D. Unpaired t-test was used in F-G and I.

### Hyperlipidemia causes severe atherosclerosis in Ccn2 deficient mice

To investigate the aorta phenotype in a hyperlipidemic setting, mice were given a single intravenous injection of 1·10^11^ viral genomes of recombinant adeno-associated virus serotype 8 harboring the D377Y gain-of-function mutant of mouse *Pcsk9* (rAAV8-D377Y-mPcsk9) downstream the hepatocyte-specific *ApoEHCR-hAAT* promoter(21). Mice were challenged with *ad libitum* western type diet (WTD) (21% fat, 0.21% cholesterol) from the day of rAAV8-D377Y-mPcsk9 injection for 24 weeks until euthanasia (**Figure 3A**). In the experimental period, *Ccn2*^*Δ/Δ*^ mice had a modest reduction in weight gain (**Figure 3B**) and plasma cholesterol (**Figure 3C**) (not statistically significant in females). After 24 weeks of hyperlipidemia, *Ccn2*^*Δ/Δ*^ aortas were severely enlarged and whitened (**Figure 3D**) with a 3-5-fold increase in relative mass (**Figure 3E**). The enlargement appeared to hinder natural alignment with the spine, and a resulting kink of the thoracic aortas was consistently observed (**Figure 3F**). Relative heart and kidney mass were increased in *Ccn2*^*Δ/Δ*^ mice (not statistically significant in males) (**Figure 3G-H**), and relative spleen mass tended to be higher in *Ccn2*^*Δ/Δ*^ mice (**Figure 3I**).

**Figure 3.**
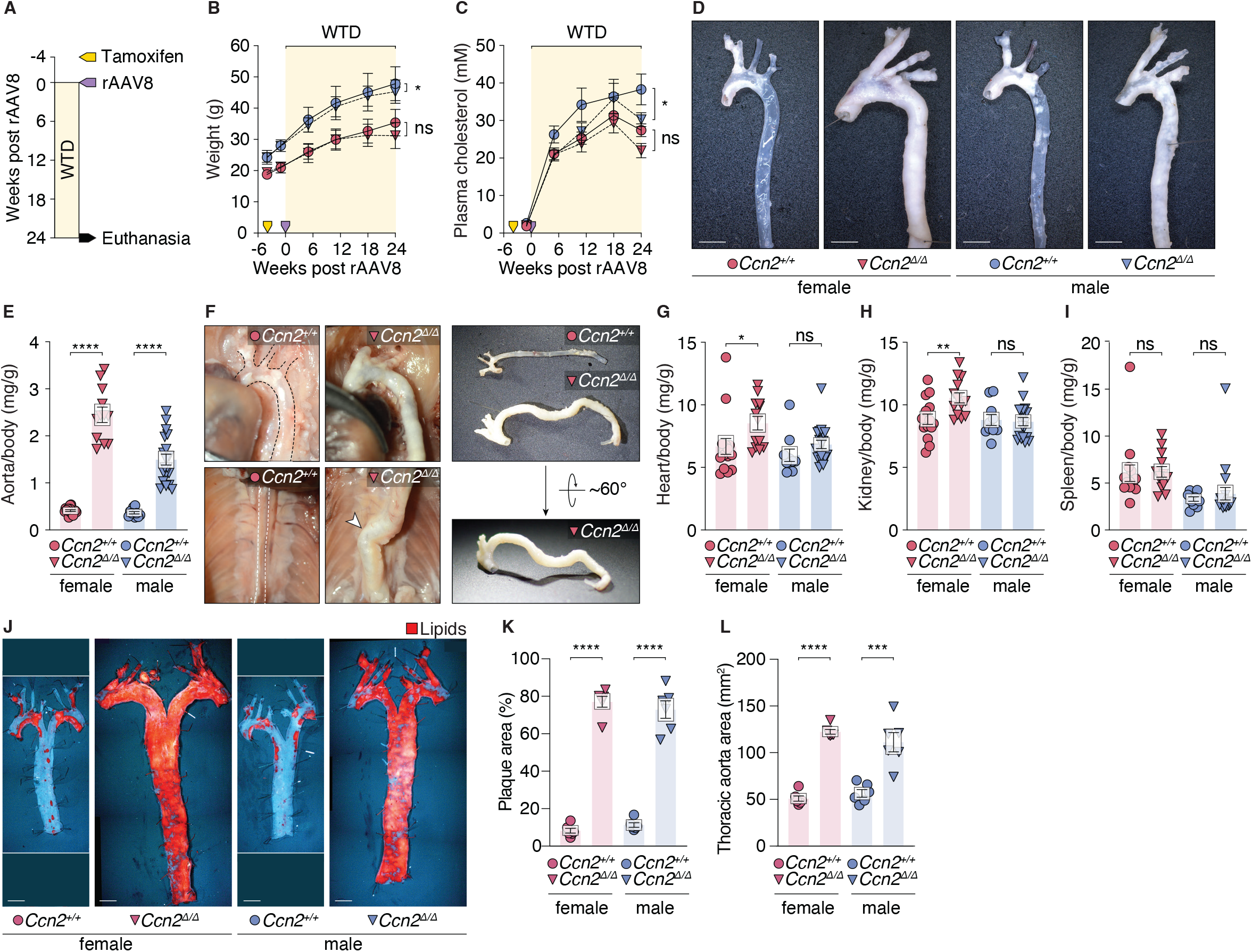
Hyperlipidemia causes severe atherosclerosis in Ccn2 deficient mice. **A**. Study design. **B-C**. Mouse weight (B) and plasma cholesterol (C) through the course of the experiment. Yellow arrowhead indicates timing of tamoxifen-treatment. Purple arrowhead indicates timing of rAAV8-D377Y-mPcsk9 injection. **D**. Representative images of aortas after 24 weeks of hyperlipidemia. **E**. Relative mass of thoracic aortas after 24 weeks of hyperlipidemia. **F**. Representative images of aortas *in situ* and *ex vivo*. While aortas of *Ccn2*^*+/+*^ mice are aligned with the spine, the enlarged aortas of *Ccn2*^*Δ/Δ*^ mice hindered natural alignment with the spine and resulted in a consistently observed kink (white arrowhead) of the thoracic aorta. **G-I**. Relative mass of heart (G), kidney (H) and spleen (I) after 24 weeks of hyperlipidemia. **J**. Oil Red O staining of *en face* prepared thoracic aortas after 24 weeks of hyperlipidemia. Scalebars = 2 mm. **K**. Percentage of thoracic aortic area stained positive for lipid. **L**. Total thoracic aortic area. Plots in the figure show mean ± SEM, except for error bars in B, which represent standard deviation. *: *p* ≤ 0.05, **: *p* ≤ 0.01, ***: *p* ≤ 0.001, ****: *p* ≤ 0.0001, ns: non-significant. Unpaired t-test was used in G-I and K-L. Two-way ANOVA was used in B-C.

Oil Red O staining of *en face*-prepared thoracic aortas showed that approximately 75% of the aortic area of *Ccn2*^*Δ/Δ*^ mice stained positive for lipid in contrast to 10% in aortas from *Ccn2*^*+/+*^ mice (**Figure 3J-K**). Moreover, the thoracic aortic area was increased by 1.75 fold (**Figure 3L**). Neointima had formed in *Ccn2*^*Δ/Δ*^ aortas and the CSA of the underlying media was significantly increased (**Figure 4A-C**). Most neointimal cells were associated with lipid and could be classified as either Galectin-3 (Lgals3)-positive foam cells, or Lgals3-negative cells (**Figure 4D-E**). Negligible lipid was detected in sections of *Ccn2*^*+/+*^ descending thoracic aortas (**Figure 4F**).

**Figure 4.**
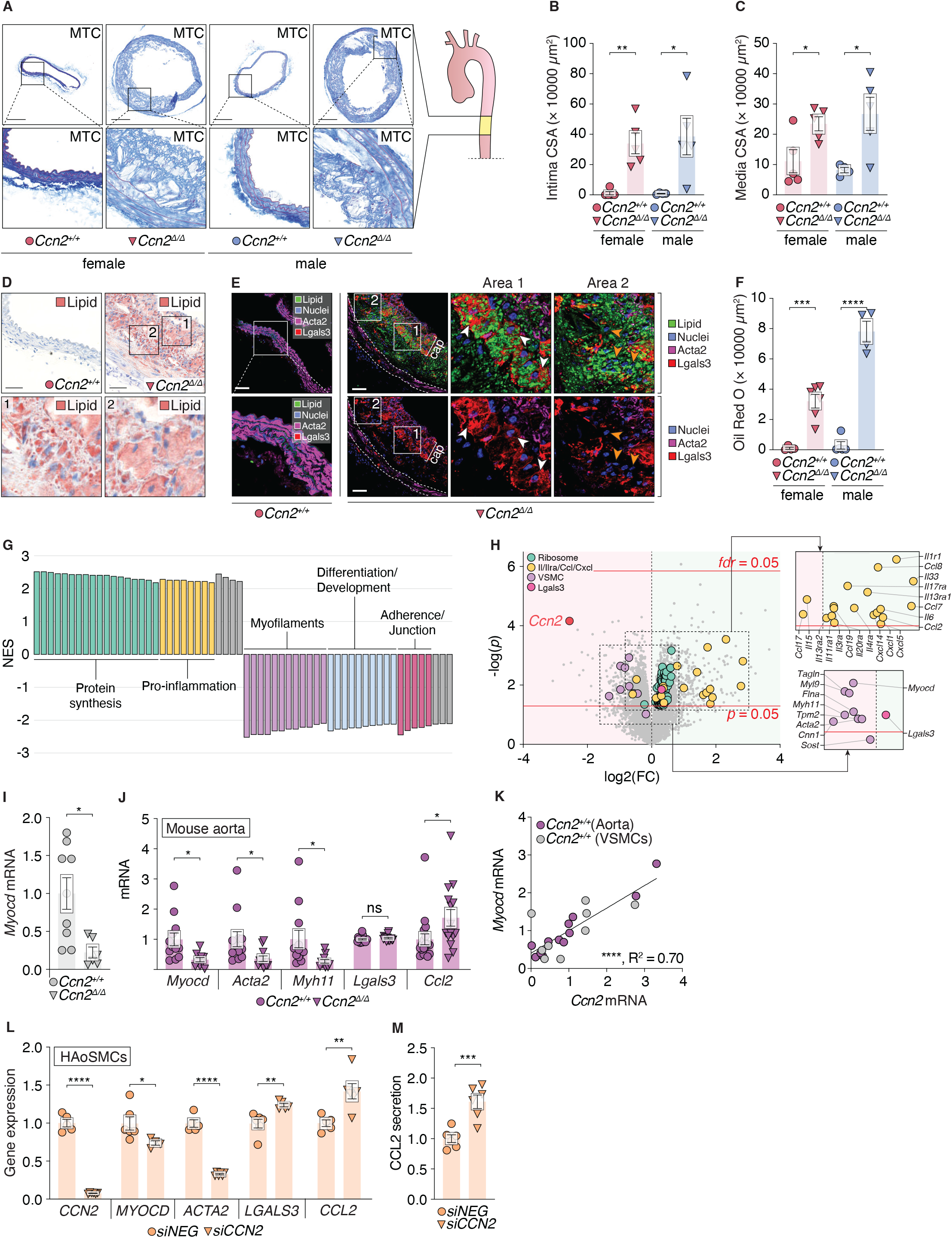
Ccn2-deficiency causes dedifferentiation of SMC. **A**. Representative cross-sections of descending thoracic aortas (indicated in yellow) stained by MTC. Scalebars = 250 *μ*m. **B-C**. Quantitation of intimal (B) and medial (C) CSA. **D**. Representative Oil Red O-staining of cross-sections shown in A. Scalebars = 50 *μ*m. **E**. Immunofluorescence co-staining of *Ccn2*^*+/+*^ and *Ccn2*^*Δ/Δ*^ aortas of Acta2 (staining was performed with an Alexa Fluor 647-conjugated secondary antibody but shown in magenta), Lgals3 (red), lipids (green) and nuclei (blue). In the bottom row images of *Ccn2*^*Δ/Δ*^ aortas, the green channel is omitted for clarity. White arrowheads point to Lgals3-positive cells. Orange arrowheads point to Lgals3-negative cells. Scalebars = 50 *μ*m. **F**. Quantitation of Oil Red O staining of sections shown in D. **G**. GSEA of microarray data from *Ccn2*^*+/+*^ and *Ccn2*^*Δ/Δ*^ SMCs. The normalized enrichment scores (NES) of the 30 top enriched gene sets with NES > 0 and NES < 0, respectively, are shown and sorted by common terms (refer to Supplementary Table 1 for identities of individual gene sets). **H**. Volcano plot showing regulation of all genes (grey dots) in *Ccn2*^*Δ/Δ*^ SMCs. Ribosomal subunits (*p* ≤ 0.05) are shown magnified in green, cytokines/ chemokines and receptors (*p* ≤ 0.05) are shown magnified in yellow (identities are shown in the top parallel-shifted rectangle), the *a priori* selected panel of eight SMC differentiation markers are shown magnified in magenta (identifies are shown in the bottom parallel-shifted rectangle), Lgals3 is shown magnified in pink, and Ccn2 is shown magnified in red. **I**. Validation of reduced *Myocd* in *Ccn2*^*Δ/Δ*^ SMCs by qPCR. **J**. Expression of selected transcripts in *Ccn2*^*Δ/Δ*^ and *Ccn2*^*+/+*^ aortas. **K**. Correlation between *Ccn2* and *Myocd* in *Ccn2*^*+/ +*^ aortas and *Ccn2*^*+/+*^ SMC normalized to *Gapdh* and then the mean value for each sample type. **L**. Expression of selected transcripts in *CCN2*-silenced HAoSMCs. **M**. CCL2 level in conditioned media of HAoSMCs. Data in L-M are from three independent experiments. Barplots in the figure show mean ± SEM. *: *p* ≤ 0.05, **: *p* ≤ 0.01, ***: *p* ≤ 0.001, ****: *p* ≤ 0.0001, ns: non-significant. Unpaired t-test was used in B, C, F, I, J, L and M and I. Microarray data in H were analyzed by unpaired t-test for each gene followed by *fdr* correction for multiple testing. Data in K were analyzed by linear regression analysis.

### Ccn2-deficiency causes dedifferentiation of SMCs

As aortas of normolipidemic *Ccn2*^*Δ/Δ*^ mice had a severe phenotype without a marked presence of tissue leukocytes (data not shown) and since SMCs appeared to be the predominant source of Ccn2 in the artery wall, we hypothesized that SMCs are the principal cells driving the vascular phenotype. To further investigate this, we isolated RNA from primary cultures of aorta-derived SMCs from normolipidemic *Ccn2*^*+/+*^ and *Ccn2*^*Δ/Δ*^ mice, and evaluated differential gene expression by microarray analysis. Genes representing GO-terms associated with protein synthesis and pro-inflammation were increased in *Ccn2*^*Δ/Δ*^ SMCs, while genes representing GO-terms associated with myofilaments, differentiation/development and adherence/junction were reduced (**Figure 4G**). One gene, *Homeobox Protein HB24* (*Hlx*), was significantly regulated (up) after correcting for multiple testing. We next looked up *Myocd* and a panel of eight genes (*Acta2, Cnn1, Myl9, Myh11, Tagln, Tpm2, Sost* and *Flna*) previously identified by single cell RNA sequencing (ssRNAseq) to strongly correlate with differentiated contractile SMCs(14), and found all but *Sost* to be downregulated in *Ccn2*^*Δ/Δ*^ SMCs (**Figure 4H**). *Myocd* was reduced by 75% in *Ccn2*^*Δ/Δ*^ SMCs (**Figure 4I**). Further, the expression of *Lgals3* (suggested marker of SMC modulation(14)), most cytokines, chemokines and receptors of these, as well as genes encoding ribosomal subunits were nominally increased (**Figure 4H**). Most of these effects were validated *in vivo* (**Figure 4J**). Moreover, in *Ccn2*^*+/+*^ aortas and *Ccn2*^*+/+*^ SMCs, *Ccn2* displayed strong correlation with *Myocd* (**Figure 4K**). Finally, *CCN2* silencing in primary human aortic smooth muscle cells (HAoSMCs) resulted in significant reduction of *MYOCD* and *ACTA2*, a significant increase in *LGALS3* and *CCL2* (**Figure 4L**) and elevated secretion of CCL2 (**Figure 4M**).

## DISCUSSION

We show that *CCN2* is one of the most abundant transcripts in non-diseased human arteries, and that CCN2 has a role in maintaining vascular integrity and protection against atherosclerosis. Further, we highlight that relative *CCN2* expression is higher in artery tissues as compared to all other tissues from the GTEx portal(19). Elevated relative expression of *CCN2* in artery tissue is also the case in mice - at least in embryos - where the vasculature is among few tissues with strong *Ccn2* expression(20). Nevertheless, the role of CCN2 in the vasculature has received little attention as the vast majority of reports have focused on the involvement of CCN2 in pathophysiological contexts of other tissues(16).

Our data imply that the observed vascular phenotype is driven by a primary effect on SMCs. First, SMCs are the principal source of *CCN2* in mouse and human arteries; second, isolated mouse *Ccn2*^*Δ/Δ*^ SMCs downregulate *Myocd* and modulate toward a less differentiated, pro-inflammatory, synthetic phenotype, as verified in HAoSMCs and *in vivo*; and finally, the aorta phenotype of normolipidemic mice occurs in the absence of detectable leukocytes. Therefore, it is conceivable that atherogenic processes, *i*.*e*. lipid accumulation and recruitment of myeloid cells, are aggravated as a consequence of a structurally altered aorta with dysfunctional SMCs.

Interestingly, in non-vascular contexts, CCN2 has previously been reported to be important for differentiation into ACTA2^+^ myofibroblasts - the principal effector cell in fibrosis. Upon kidney injury, tubular epithelial cells require CCN2 to transdifferentiate into contractile myofibroblasts that induce fibrosis(16, 22); and CCN2 is required for differentiation of dermal progenitor cells into myofibroblast-like cells(23). Moreover, CCN2 drives early differentiation of myoblasts in mice while it inhibits terminal differentiation of skeletal myoblasts(24).

In conclusion, we find that CCN2 plays a critical protective role in artery tissues, and is required to maintain SMCs in a differentiated, contractile phenotype. To this end, our data argue caution in using anti-CCN2-based therapy.

## METHODS

### In silico analysis

Relative transcript levels of *CCN2* (expressed as transcripts per million, TPM) were assessed using GTEx Portal V8(19). Of note, the GTEx Portal uses *CTGF* as gene symbol rather than *CNN2*.

### Mouse procedures

*Ccn2*^*fl/fl*^*Rosa26-CreER*^*T2*^ mice (C57BL/6J) were crossed with C57BL/6J (Taconic, Denmark) to generate *loxp*-flanked *Ccn2* exon 4 mice (*Ccn2*^*fl/fl*^) either carrying or not carrying the *Rosa26-CreER*^*T2*^ knockin. Primers used for genotyping were: *Ccn2* locus: 5’AATACCAATGCACTTGCCTGGATGG and 5’-GAAACAGCAATTACTACAACGGGAGTGG; *Rosa26* locus:

5’AAAGTCGCTCTGAGTTGTTAT, 5’GGAGCGGGAGAAATGGATATG and 5’CCTGATCCTGGCAATTTCG.

At 8 weeks of age, mice were intraperitoneally injected with 1 mg tamoxifen (T5648, Sigma Aldrich) in 100 *μ*L corn oil (C8267, Sigma Aldrich) each day for four consecutive days. Six weeks post tamoxifen treatment, mice were euthanized by CO^2^/O^2^ inhalation and tissues were either snap frozen in liquid nitrogen or immersed in 4% formaldehyde in phosphate-buffered saline (PBS) for 24 hours and then transferred to PBS supplemented with 0.05% sodium azide.

Hyperlipidemia was induced by a single intravenous injection of 1·10^11^ viral genomes of rAAV8-D377Y-mPcsk9 produced at University of North Carolina Vector Core (Chapel Hill, NC, USA) from pAAV/ApoEHCR-hAAT-D377Y-mPcsk9-BGHpA in combination with western type diet (WTD, 21% fat, 0.21% cholesterol) (D12079B, Research Diets Inc.) for 24 weeks beginning at the day of rAAV8-D377Y-mPcsk9 injection. Every six weeks, mice were weighed and blood was harvested from the facial vein to monitor plasma cholesterol (CHOL2, Roche Diagnostics), and six hour fasting blood glucose was measured four weeks into the experiment and four weeks prior to euthanasia (tail tip bleed readings by Accu-Chek, Roche). After 24 weeks of hyperlipidemia, mice were euthanized by intraperitoneal injection of pentobarbital (250 mg/kg) and lidocain (20 mg/kg), and then perfused at 100 mmHg with PBS for 30 sec followed by 4% formaldehyde in PBS for 5 min through the left ventricle using the cut right atrium as route of drainage. Mice were then immersed in 4% formaldehyde in PBS for six hours and stored in PBS supplemented with 0.05% sodium azide at 4°C.

Mouse group allocation was random (littermates were used for all experiments and *Ccn2*^*Δ/Δ*^ and *Ccn2*^*+/+*^ shared cage), concealed, and all experimental procedures and analyses were performed blinded to the investigator. However, the severity of the *Ccn2*^*Δ/Δ*^ phenotype was self-revealing in most morphometric and histological analyses. All mice from litters of all synchronized breeding were included in experiments and no mice were excluded from experiments other than death. A total of one normolipidemic mouse (male *Ccn2*^*Δ/Δ*^) and three hyperlipidemic mice (1 female *Ccn2*^*+/+*^, 1 female *Ccn2*^*Δ/Δ*^, and 1 male *Ccn2*^*Δ/Δ*^) died from unknown causes.

All procedures were performed in accordance with protocols approved by the local authorities (the Danish Animal Experiments Inspectorate) and conform to the guidelines from Directive 2010/63/EU of the European Parliament on the protection of animals used for scientific purposes.

### Transcriptomics

RNA was extracted from snap frozen tissue or cells using NORGEN Total RNA Purification Kit (37500, Norgen Biotek Corp.) and cDNA was made by High-Capacity cDNA Reverse Transcription Kit (4368814, Thermo Fisher Scientific). Quantitative PCR was performed using SYBR™ Green PCR Master Mix (4364344, Thermo Fisher Scientific) and the ViiA7 Real-Time PCR system (Thermo Fisher Scientific). Primer pairs used for mouse transcripts were: *Ccn2*: 5’GAGGAAAACATTAAGAAGGGC and 5’CAGGCACAGGTCTTGATGAAC; *Acta2*: 5’CCTTCGTGACTACTGCCGAG and 5’ATAGGTGGTTTCGTGGATGC; *Myocd*: 5’CCTGCCTGCTGGGAGTTAAT and 5’GCCGTAACTGTAAGACTGATCG; *Ccl2*: 5’ CACTCACCTGCTGCTACTCA and 5’ GCTTGGTGACAAAAACTACAGC; *Gapdh*: 5’GGCTGCCCAGAACATCAT and 5’CGGACACATTGGGGGTAG. Primer pairs used for human transcripts were: *CCN2*: 5’CATCTTCGGTGGTACGGTGT and 5’TTCCAGTCGGTAAGCCGC; *ACTA2*: 5’CCAGAGCCATTGTCACACAC and 5’CAGCCAAGCACTGTCAGG; *MYOCD*: 5’ AAAAAGCCCAAGGACCCCAA and 5’GTCCATAGGTGGAGGGGACT; *CCL2*: 5’ TCCCAAAGAAGCTGTGATCTTCA and 5’ TCTGGGGAAAGCTAGGGGAA; *GAPDH*: 5’CAGCGACACCCACTCCTC and 5’TGAGGTCCACCACCCTGT. *Gapdh/GAPDH* was used as reference gene.

For microarray analysis, background correction, normalization, and gene expression index calculation of probe intensities were done in Transcriptome Analysis Console (TAC) software (Thermo Fisher Scientific) using the robust multi-array average (rma) method. Probes representing non-expressed genes as defined by TAC were removed from further analysis. All subsequent calculations were performed using the open source R-environment (R version 3.6.1). First, the normalized probe expression matrix was collapsed by gene symbol using maximal probe intensity, thus providing one gene expression value per sample per gene. Next, the experiment series associated batch effects were removed/reduced by COMBAT batch removal using the ComBat function from the *sva* r-package. From this matrix we selected the 2000 most variable genes and visualized gene expression levels using the R-package *ComplexHeatmap*^25^. Differential gene expression analysis between *Ccn2*^*Δ/Δ*^ and *Ccn2*^*+/+*^ mice was conducted on the gene expression measured by the microarrays using an unpaired limma t-test embedded in the *limma* R-package(25). Genes with *fdr* ≤ 0.05 were considered as being significantly differentially expressed between *Ccn2*^*Δ/Δ*^ and *Ccn2*^*+/+*^ mice. Gene set enrichment analysis (GSEA) was performed using the *fgsea* R-package to determine if a given set of curated genes/pathways showed statistically significant, concordant differences between *Ccn2*^*Δ/Δ*^ and *Ccn2*^*+/+*^ mice. The GSEA was run on pre-ranked individual expressed genes using the log2 fold change as ranking metric. The collection of mouse gene ontology (GO) defined gene sets were used as input for the analysis. Next, an enrichment score was calculated for each of these *a priori* defined gene sets. In our analysis, GSEA was conducted using 1000 permutations, and minimum and maximum gene sets size was set to 15 and 500, respectively. Gene sets with *fdr* ≤ 0.05 were considered as being statistically significantly enriched.

### Proteomics

Snap frozen tissues were homogenized using a TissueLyser system (Qiagen) with stainless steel beads (Qiagen) in a lysis buffer (100 mM DTT, 5% sodium deoxycholate, 1% β-octylglucoside, 20 mM Tris, pH 8.8, supplemented with complete, Mini, EDTA-free Protease Inhibitor Cocktail Tablets and PhosSTOP (Roche)). Proteins were then denatured (99°C for 20 min, 80°C for 100 min), alkylated (150 mM iodoacetamide for 30 min at room temperature), precipitated (−20°C acetone for 1 hour), resolubilized (8M urea for 30 min at room temperature), and trypsinized (0.5 *μ*g/*μ*l trypsin (Promega) in 50 mM ammonium bicarbonate, 1M urea at 37°C overnight). After acidification, tryptic peptides were purified on custom-made Poros R2/R3 (Thermo Scientific) micro-columns, dried, reconstituted, and finally resuspended in 2% acetonitrile (ACN), 0.1% formic acid. The concentration of tryptic peptides was determined (Pierce BCA Protein Assay Kit, Thermo Fisher Scientific) and normalized across samples.

For targeted MRM analysis, heavy isotope-labeled standard peptides (JPT Peptide Technologies) were added to each sample aiming at a 0.1-10 peak area ratio between each endogenous peptide assayed and its corresponding heavy peptide. Each sample (1 *μ*g endogenous tryptic peptides) was run on an Easy-nLC II NanoLC system using a C18 trapping column (length 2 cm; internal diameter 100 *μ*m) for desalting and a C18 analytical column (length 10 cm, internal diameter 75 *μ*m) for peptide separation (Thermo Scientific). Peptides were eluted with a three-step 45 min gradient of 0.1% formic acid in ACN at a flow rate of 300 nl/min. Peptides were ionized using a Nanospray Flex ion source (Thermo Scientific) and analyzed on a TSQ Vantage triple quadruple mass spectrometer (Thermo Scientific) in MRM mode. MRM raw files were processed using Pinpoint 1.3 (Thermo Scientific). The peak area ratio between endogenous and heavy isotope labeled spiked peptide was used for data analysis. The peak area ratio for Ccn2 (target peptide: TTTLPVEFK) was normalized to Gapdh (target peptides: GAAQNIIPASTGAAK and LISWYDNEYGYSNR) within the same sample (**Figure 2E**).

### Whole aorta analysis

Formalin-fixed aortas from the beginning of the aortic arch to the diaphragm were dissected, cut open longitudinally and stained with a 2:1 mix of 0.5% Oil Red O in isopropanol (O1391, Sigma Aldrich) and deionized water for 10 min at 37°C, briefly washed in isopropanol (I9516, Sigma Aldrich) and transferred to deionized water. Aortas were pinned to a black rubber material, immersed in PBS and images were generated using a Zeiss AxioCam ERc 5s camera attached to a Zeiss Stemi 2000-C microscope. Thoracic aortic area and percentage of thoracic aorta stained by Oil Red O, was determined using an automated macro in Fiji (ImageJ).

### Tissue preparation and histology

Human internal thoracic arteries and carotid plaques were obtained from coronary artery bypass grafting and carotid endarterectomy procedures, respectively, performed at Odense University Hospital. The only parameter used to select human specimens was that randomly chosen samples had to be morphologically acceptable. All donors had given written consent, and the study was performed in accordance with protocols approved by the local ethics committee (S-20100044) and conformed to the principles of the Declaration of Helsinki. Tissue was fixed in 4% formaldehyde in PBS for 24 hours, embedded in paraffin, sectioned at 5 *μ*m thickness, and deparaffinized prior to histological staining.

Formalin fixed mouse aortas were cryoprotected in sucrose (25% in PBS for 24 hours, then 50% in PBS for 24 hours), embedded in optimal cutting temperature (OCT) compound (Tissue-Tek, Sakura), and sectioned at 5 *μ*m thickness.

For Masson trichome (MTC) staining, sections were airdried, and exposed to: Papaniculaus 1 (6 min), running tap water (8 min), 1.2% picric acid (5 min), tap water (1 min), 1% Biebrick scarlet (10 min), 1% phosphor wolfram acid (10 min), 2.5% methyl blue (2 min), acetic acid (30 sec), 1% phosphor wolfram acid (5 min), and acetic acid (3 min). Slides were then dehydrated and coverslips were mounted using Aquatex (Sigma Aldrich). For Oil Red O staining, sections were exposed to: 7% ethanol (1 min), Oil Red O (2.5 min), 100% ethanol (1 min), tap water (1 min), Mayer’s Hematoxylin (2 min), tap water (1 min), 0.3% sodium carbonate (10 sec), running tap water (1 min). Coverslips were then mounted using Aquatex (Sigma Aldrich).

### In situ hybridization

*In situ* hybridization (ISH) was performed with RNAscope® 2.5 VS Reagent Kit (323250, Advanced Cell Diagnostics, Newark, CA) on a Ventana Discovery Ultra platform (Roche Diagnostics, Penzberg, Germany). Sections were exposed to 24 hours antigen retrieval and 16 min protease treatment. The probes applied were specific for human *CCN2* mRNA (cat no. 560589), and for *dapB* (*4-hydroxy-tetrahydrodipicolinate reductase* from *Escherichia coli*) as negative control (cat no. 312039). Signal was detected using Fast Red detection kit (8127166001, Roche Diagnostics, Penzberg, Germany).

### Immunohistochemistry and immunofluorescence

For Cd45 immunohistochemistry, sections were airdried, washed in Tris-buffered saline (TBS) (3 × 5 min) and demasked in citrate buffer (10 mM citric acid monohydrate, pH 6.0) at 97°C (20 min). After reaching room temperature, sections were blocked in 1.66% H^2^O^2^ in TBS (10 min), washed in TBS (3 × 5 min) and blocked in 1% BSA (Sigma Aldrich) in TBST (0.1% Tween-20 in TBS). Sections were then incubated with rat anti-mouse Cd45 antibody (55039, BD Pharmingen) (1:100 in 1% BSA in TBST) overnight at 4°C, washed in TBST (3 × 5 min), incubated with HRP-conjugated rabbit anti-rat IgG (P0450, Dako) (1:1000 in 1% BSA in TBST), and washed in TBST (3 × 5 min). DAB solution (K3568, Dako) was applied (14 min) and sections were washed in tap water (5 min) and counterstained with Mayer’s Hematoxylin (30 sec) and mounted using Aquatex (Sigma Aldrich).

For co-staining of Acta2, Lgals3, nuclei and lipids, sections were airdried, washed in deionized water (1 min) and PBS (3 min), blocked in 5% normal goat serum (NGS) (ab7481, Abcam) in PBS (30 min), incubated with rabbit anti-mouse Acta2 antibody (ab5694, Abcam) (1:300 in 5% NGS in PBS) and rat anti-mouse Lgals3 antibody (M3/38, Cedarlane) (1:600 in 5% NGS in PBS) (2 hours), washed in PBS (3 × 3 min), incubated with Alexa Fluor 647-conjugated goat anti-rabbit IgG (A21244, Invitrogen) (1:300 in 5% NGS in PBS), Alexa Fluor 555-conjugated goat anti-rat IgG (A21434, Invitrogen) (1:300 in 5% NGS in PBS), BODIPY (493/503) (D3922, Invitrogen) (10 mg/ ml in DMSO) (1:600 in 5% NGS in PBS) (1 hour), and washed in PBS (3 × 3 min). Coverslips were then mounted with SlowFade Gold Antifade Mountant with DAPI (S36942, Thermo Fisher Scientific). Images were recorded by a Hamamatsu model 2 OHT scanner and immunofluorescence images were recorded with an Olympus FV1000MPE upright confocal laser scanning confocal microscope (excitation lasers 405 nm, 488 nm, 559 nm and 635 nm) using the FV-ASW (version 4.2C) software.

ACTA2, MYH11 and CD68 immunohistochemistry of human tissues was performed at the Department of Pathology, Odense University Hospital, using primary antibodies BS66 (BSH-7459-100, Nordic Biosite, 1:1000), SMMS-1 (M3558, Dako, 1:100) and PG-M1 (M0876, Dako, 1:50), respectively, utilizing the full-automated OptiView DAB IHC Detection Kit (760-700). Images were recorded by a Hamamatsu model 2 OHT scanner.

### SMC cultivation

*Ccn2*^*Δ/Δ*^ and *Ccn2*^*+/+*^ mice were euthanized by cervical dislocation six weeks post tamoxifen treatment. The right atrium was cut and mice were flushed via the left ventricle using sterile PBS (70011044, Thermo Fisher Scientific). The descending thoracic aortas were dissected free from perivascular tissue while in normal growth media (DMEM supplemented with 20% fetal bovine serum (FBS) (10270106, Life Technologies), 1% Penicillin-Streptomycin (15140122, Life Technologies), 1% L-Glutamine (G7513, Sigma Aldrich) containing 0.01% Amphotericin B (15290018, Thermo Fisher Scientific)). Aortas were cut into 1×1 mm pieces, placed in a 96-well plate (6267170, Buch & Holm) and digested in normal growth media containing 1.42 mg/ml Collagenase (C6885, Sigma Aldrich) for 6 hours at 37ᵒC. Cell suspensions were then centrifuged at 350 x *g* for 5 min, resuspended in normal growth media, and seeded in 24-well plates (6267168, Buch & Holm). Fourth passage cells were starved for 48 hours in normal growth media (without FBS), trypsinated (25200072, Sigma Aldrich), centrifuged for 350 x *g* for 5 min, and stored at -80ᵒC until use.

HAoSMCs (C-12533, PromoCell) were grown in Smooth Muscle Cell Growth Medium 2 (C-22062, PromoCell). At 80% confluency, third passage cells were starved for 48 hours in Basal Medium 2, phenol red-free (C-22267, PromoCell), trypsinated (25200072, Sigma Aldrich), centrifuged at 220 x *g* for 5 min, and seeded (6000 cells/cm^2^) in Smooth Muscle Cell Growth Medium 2 in 12-well plates (6267167, Buch & Holm). After 24 hours, media was removed and cells were tranfected with 50 nM ON-TARGETplus Human CTGF (1490) siRNA - SMARTpool (L-012633-01-0005, Horizon) or ON-TARGETplus Non-targeting Pool (D-001810-10-05, Horizon) in Basal Medium 2, phenol red-free using Lipofectamine™ RNAiMAX Transfection Reagent (13778075, Thermo Fisher Scientific) according to manufacturer’s instructions for 24 hours. Tranfection media was replaced by Smooth Muscle Cell Growth Medium 2, and after 24 hours, media (centrifuged at 1500 x *g* for 10 minutes) and cells (trypsinated (25200072, Sigma Aldrich) and centrifuged at 220 x *g* for 5 min) were harvested and stored at -80ᵒC until use.

CCL2 was quantified in conditioned media from HAoSMCs using Human CCL2/MCP-1 Quantikine ELISA Kit (DCP00, R&D systems) according to the manufacture’s protocol.

### Statistical analysis

Two-group comparisons of normal distributed data were performed using unpaired t-test (with Welch’s correction if variances unequal) and presented as mean ± standard error of the mean (SEM). Comparison of groups over time was performed using Two-way ANOVA and presented as mean ± SEM (Error bars in **Figure 3B** represents standard deviation to make them visible). GO enrichment analysis results presented in **Figure 2M** was performed using default settings of the DAVID Bioinformatics Resources^28,29^. Statistical approaches used to analyze microarray data are described above.

Replicas (*n*) are specified on figures or in figure legends whenever data are not plotted as individual data points. In the experiment investigating effects of Ccn2 deficiency in the setting of hyperlipidemia, a subset of mice from each group (*Ccn2*^*+/+*^ female, *Ccn2*^*Δ/Δ*^ female, *Ccn2*^*+/+*^ male, *Ccn2*^*Δ/Δ*^ male) was randomly selected for *en face*-based analyses of thoracic aortas and histology.

In graphs where data values are not absolute, individual values are normalized to the mean of the control group (or *Ccn2*^*+/+*^ females when showing both sexes) defined as 1. *p* ≤ 0.05 was the criterion for reliable differences between groups.

### Data availability statement

The data underlying this article will be shared on reasonable request to the corresponding author.

## SOURCES OF FUNDING

Odense University Hospital

## ACKNOWLEDGEMENTS

We thank Christian Enggaard, Morten Ploug Kühlmann, Anne-Sofie Madsen, Inger Nissen, Camilla Rasmussen and Pia Søndergaard Jensen for excellent technical assistance. We thank Professor Jacob Fog Bentzon (Aarhus University Hospital, Denmark) for providing pAAV/ApoEHCR-hAAT-D377Y-mPcsk9-BGHpA.

## CONFLICTS OF INTEREST

The authors have nothing to declare.

## Notes

### Competing Interest Statement

The authors have declared no competing interest.

## REFERENCES

1. Basatemur GL, Jorgensen HF, Clarke MCH, Bennett MR, Mallat Z. Vascular smooth muscle cells in atherosclerosis. Nat Rev Cardiol. 2019;16(12):727–44.

2. Yoshida K, Gowers KHC, Lee-Six H, Chandrasekharan DP, Coorens T, Maughan EF, et al. Tobacco smoking and somatic mutations in human bronchial epithelium. Nature. 2020;578(7794):266–72.

3. Bennett MR, Sinha S, Owens GK. Vascular Smooth Muscle Cells in Atherosclerosis. Circ Res. 2016;118(4):692–702.

4. Parker F. An Electron Microscopic Study of Experimental Atherosclerosis. Am J Pathol. 1960;36(1):19–53.

5. Geer JC, Mc GH, Jr., Strong JP. The fine structure of human atherosclerotic lesions. Am J Pathol. 1961;38(3):263–87.

6. Stary HC. Composition and classification of human atherosclerotic lesions. Virchows Arch A Pathol Anat Histopathol. 1992;421(4):277–90.

7. Gomez D, Owens GK. Smooth muscle cell phenotypic switching in atherosclerosis. Cardiovasc Res. 2012;95(2):156–64.

8. Chappell J, Harman JL, Narasimhan VM, Yu H, Foote K, Simons BD, et al. Extensive Proliferation of a Subset of Differentiated, yet Plastic, Medial Vascular Smooth Muscle Cells Contributes to Neointimal Formation in Mouse Injury and Atherosclerosis Models. Circ Res. 2016;119(12):1313–23.

9. Jacobsen K, Lund MB, Shim J, Gunnersen S, Fuchtbauer EM, Kjolby M, et al. Diverse cellular architecture of atherosclerotic plaque derives from clonal expansion of a few medial SMCs. JCI Insight. 2017;2(19).

10. Gomez D, Shankman LS, Nguyen AT, Owens GK. Detection of histone modifications at specific gene loci in single cells in histological sections. Nat Methods. 2013;10(2):171–7.

11. Shankman LS, Gomez D, Cherepanova OA, Salmon M, Alencar GF, Haskins RM, et al. KLF4-dependent phenotypic modulation of smooth muscle cells has a key role in atherosclerotic plaque pathogenesis. Nat Med. 2015;21(6):628–37.

12. Albarran-Juarez J, Kaur H, Grimm M, Offermanns S, Wettschureck N. Lineage tracing of cells involved in atherosclerosis. Atherosclerosis. 2016;251:445–53.

13. Misra A, Feng Z, Chandran RR, Kabir I, Rotllan N, Aryal B, et al. Integrin beta3 regulates clonality and fate of smooth muscle-derived atherosclerotic plaque cells. Nat Commun. 2018;9(1):2073.

14. Wirka RC, Wagh D, Paik DT, Pjanic M, Nguyen T, Miller CL, et al. Atheroprotective roles of smooth muscle cell phenotypic modulation and the TCF21 disease gene as revealed by single-cell analysis. Nat Med. 2019;25(8):1280–9.

15. Perbal B, Tweedie S, Bruford E. The official unified nomenclature adopted by the HGNC calls for the use of the acronyms, CCN1-6, and discontinuation in the use of CYR61, CTGF, NOV and WISP 1-3 respectively. J Cell Commun Signal. 2018;12(4):625–9.

16. Ramazani Y, Knops N, Elmonem MA, Nguyen TQ, Arcolino FO, van den Heuvel L, et al. Connective tissue growth factor (CTGF) from basics to clinics. Matrix Biol. 2018;68-69:44–66.

17. Rodrigues-Diez RR, Tejera-Munoz A, Esteban V, Steffensen LB, Rodrigues-Diez R, Orejudo M, et al. CCN2 (Cellular Communication Network Factor 2) Deletion Alters Vascular Integrity and Function Predisposing to Aneurysm Formation. Hypertension. 2022;79(3):e42–e55.

18. Wang Y, Liu X, Xu Q, Xu W, Zhou X, Lin Z. CCN2 deficiency in smooth muscle cells triggers cell reprogramming and aggravates aneurysm development. JCI Insight. 2023;8(1).

19. Consortium GT, Laboratory DA, Coordinating Center -Analysis Working G, Statistical Methods groups-Analysis Working G, Enhancing Gg, Fund NIHC, et al. Genetic effects on gene expression across human tissues. Nature. 2017;550(7675):204–13.

20. Ivkovic S, Yoon BS, Popoff SN, Safadi FF, Libuda DE, Stephenson RC, et al. Connective tissue growth factor coordinates chondrogenesis and angiogenesis during skeletal development. Development. 2003;130(12):2779–91.

21. Bjorklund MM, Hollensen AK, Hagensen MK, Dagnaes-Hansen F, Christoffersen C, Mikkelsen JG, et al. Induction of atherosclerosis in mice and hamsters without germline genetic engineering. Circ Res. 2014;114(11):1684–9.

22. Vanhove T, Goldschmeding R, Kuypers D. Kidney Fibrosis: Origins and Interventions. Transplantation. 2017;101(4):713–26.

23. Liu S, Herault Y, Pavlovic G, Leask A. Skin progenitor cells contribute to bleomycin-induced skin fibrosis. Arthritis Rheumatol. 2014;66(3):707–13.

24. Nishida T, Kubota S, Aoyama E, Janune D, Lyons KM, Takigawa M. CCN family protein 2 (CCN2) promotes the early differentiation, but inhibits the terminal differentiation of skeletal myoblasts. J Biochem. 2015;157(2):91–100.

25. Ritchie ME, Phipson B, Wu D, Hu Y, Law CW, Shi W, et al. limma powers differential expression analyses for RNA-sequencing and microarray studies. Nucleic Acids Res. 2015;43(7):e47.

